# Cross-modal restoration of juvenile-like ocular dominance plasticity after increasing GABAergic inhibition

**DOI:** 10.1101/416248

**Authors:** Manuel Teichert, Marcel Isstas, Franziska Wieske, Christine Winter, Jürgen Bolz

## Abstract

In juvenile and “young adult” mice monocular deprivation (MD) shifts the ocular dominance (OD) of binocular neurons in the primary visual cortex (V1) away from the deprived eye. However, OD plasticity is completely absent in mice older than 110 days, but can be reactivated by treatments which decrease GABA levels in V1. Typically, these OD shifts can be prevented by increasing GABAergic transmission with diazepam. We could recently demonstrate that both bilateral whisker and auditory deprivation (WD, AD), can also restore OD plasticity in mice older than 110 days, since MD for 7 days in WD mice caused a potentiation of V1 input through the ipsilateral (open) eye, the characteristic feature of OD plasticity of “young adult” mice. Here we examined whether WD for 7 days also decreases GABA levels. For this, we performed post mortem HPLC analysis of V1 tissue. Indeed, we found that WD significantly decreased GABA levels in V1. Surprisingly, enhancing GABAergic inhibition by diazepam did not abolish OD shifts in WD mice, as revealed by repeated intrinsic signal imaging. On the contrary, this treatment led to a depression of V1 input through the previously closed contralateral eye, the characteristic signature of OD plasticity in juvenile mice during the critical period. Interestingly, the same result was obtained after AD. Taken together, these results suggest that cross-modally restored OD plasticity does not only depend on reduction of GABA levels in V1, but also requires other, so far unknown mechanisms.

## Introduction

The primary visual cortex (V1) of rodents is dominated by the input from the contralateral eye (Drager, 1975, 1978). However, MD for a few days can shift the ocular dominance (OD) away from the closed eye (Wiesel and Hubel, 1963; Gordon and Stryker, 1996). In mice, this effect appears to be strongest between 28 and 32 days of age, the peak of the so called “critical period” (Gordon and Stryker, 1996). MD during this period leads to a reduction of V1 inputs from the previously closed eye, the characteristic signature of “juvenile” OD plasticity (Gordon and Stryker, 1996; Hofer et al., 2006; Kaneko et al., 2008). In young adult mice around 60 days of age, however, the mechanism, that leads to OD shifts, changes, as in these animals MD causes a potentiation of V1 responses to the input through the open eye (Sawtell et al., 2003; Sato and Stryker, 2008; Ranson et al., 2012). These changes are broadly referred to as “adult” OD plasticity. However, OD plasticity shows an age dependent decline and is completely absent in mice older than 110 days (Lehmann and Lowel, 2008). It has been suggested that the cortical inhibitory tone, which increases during aging, triggers closing of OD plasticity in fully adult mice (Hensch, 2005; Espinosa and Stryker, 2012; Levelt and Hubener, 2012). Indeed, interventions that can restore OD-plasticity in these older mice typically decrease GABA levels in V1 and thus alter the balance between excitation and inhibition (E/I balance) (Hubener and Bonhoeffer, 2014). These OD shifts, however, can be prevented or at least markedly reduced by artificially strengthening GABAergic inhibition with diazepam, a positive allosteric modulator of GABA_A_ receptors (Maya-Vetencourt et al., 2008; Spolidoro et al., 2011; Greifzu et al., 2014; Stodieck et al., 2014), suggesting that reducing cortical inhibition is a central hub to restore cortical plasticity in adults.

It has been demonstrated that deprivation of non-visual sensory modalities, such as WD or AD, in fully adult mice lead to compensatory neuronal changes in the spared V1, which are accompanied by improved visual perception at both, the physiological and behavioral level (Teichert and Bolz, 2017b; Teichert et al., 2018b). This type of plasticity is called “cross-modal plasticity” (Bavelier and Neville, 2002; Lee and Whitt, 2015). Moreover, we could recently also show that these sensory deprivations can even restore the “adult” form of OD plasticity in V1 of older mice, since combining either WD or AD, respectively, with MD resulted in a marked increase of V1 responsiveness to open eye stimulation (Teichert et al., 2018a).

Here we investigated whether cross-modally induced OD plasticity is also accompanied by reductions in GABA levels in V1, and, if so, whether OD shifts can be prevented by increasing the inhibitory tone via diazepam administration. Here we found that WD, indeed, cross-modally reduces GABA levels in the spared V1 of fully adult mice, as revealed by post-mortem HPLC analysis. These results indicate that WD decreases inhibition in V1. Using repeated intrinsic signal imaging we found that WD in saline treated control animals restored the “adult” form of OD plasticity, confirming our recently published finding (Teichert et al., 2018a). However, unexpectedly, when we increased the inhibitory tone in V1 by diazepam administrations, OD plasticity was not prevented. Quite the opposite, this treatment changed the signature of OD shifts from the “adult form” to the “juvenile form”, as we found a significant reduction in V1 responses evoked by visual stimulation of the previously closed contralateral eye. Interestingly, we could also show that increasing inhibition in mice after AD also induced “juvenile-like” OD changes, suggesting that this effect is a general feature of cross-modally restored OD plasticity. Moreover, these V1 input changes required NMDA receptor activation, as administration of the NMDA receptor antagonist CPP abolished “juvenile-like” OD shifts, emphasizing the pivotal role of these receptors in cross-modally induced plasticity. To the best of our knowledge, this is the first demonstration that increasing inhibition in the fully adult V1 does not abolish restoration of OD plasticity, but rather leads to a quality change of OD shifts. Taken together, our data suggest that cross-modal induction of cortical plasticity is not only a result of decreased inhibition, but also requires additional, so far unknown mechanisms.

## Experimental procedures

### Animals

C57BL/6J (Jackson labs) mice were raised in a group of 2-3 in transparent standard cages (16.5×22.5 cm) on a 12 h light/dark cycle, with food and water available *ad libitum*. Between the chronic experiments each animal was housed alone in a standard cage. The environment in the cage was minimally enriched with cotton rolls and nest material. In our mouse facility the light intensity was about 150-170 lux. As demonstrated recently, these rearing conditions are not sufficient to extend OD plasticity into adulthood (Teichert et al., 2018a). Animal housing in our institution is regularly supervised by veterinaries from the state of Thuringia, Germany. For the present study we used a total of 37 fully adult male and female mice (Postnatal day (PD) 120-240). All experimental procedures have been performed according to the German Law on the Protection of Animals and the corresponding European Communities Council Directive 2010 (2010/63/EU), and were approved by the Thüringer Landesamt für Lebensmittelsicherheit und Verbraucherschutz (Thuringia State Office for Food Safety and Consumer Protection) under the registration numbers 02-032/16 and 02-050/14. Every effort was made to minimize the number of animals used and their suffering.

### High performance liquid chromatography (HPLC)

WD for subsequent HPLC analysis was performed as described previously (Teichert et al., 2018b; Teichert et al., 2018a) (see below). Micropunches of V1 were taken from 1 mm brain slices at −3.28 from Bregma from control (n=5) and WD mice (n=6, after 7 days) and homogenized by ultrasonication in 20 vol of 0.1 N perchloric acid at 4 °C immediately after collection. A total of 100 ml of the homogenate was added to equal volumes of 1 N sodium hydroxide for measurement of protein content. The remaining homogenate was centrifuged at 17 000 g and 4 °C for 10 min. Glutamate and GABA levels were determined using methods described previously (Winter et al., 2009). Briefly, amino acids were precolumn-derivatized with o-phthalaldehyde-2-mercaptoethanol using a refrigerated autoinjector and then separated on a HPLC column (ProntoSil C18 ace-EPS) at a flow rate of 0.6 ml/min and a column temperature of 40 °C. The mobile phase was 50 mM sodium acetate (pH 5.7) in a linear gradient from 5% to 21% acetonitrile. Derivatized amino acids were detected by their fluorescence at 450 nm after excitation at 330 nm.

### Optical imaging of intrinsic signals

#### Mouse preparation for optical imaging

Intrinsic signal imaging experiments were performed in a total of 26 animals. In order to measure visually evoked activity of V1 we used Fourier based periodic intrinsic signal imaging (Kalatsky and Stryker, 2003; Isstas et al., 2017; Teichert et al., 2018a). Animals were initially anesthetized with 4% isoflurane in a mixture of 1:1 O_2_/N_2_O and placed on a heating blanket for maintaining body temperature (37.5°C). Subsequently, mice received injections of chlorprothixene (20 µg/mouse i.m.) and carprofene (5 mg/kg, s.c.). The animal was fixed in a stereotaxic frame and we removed the skin of the left hemisphere to expose the visual cortex. The exposed area was covered with 2.5% agarose in saline and sealed with a standard microscope glass coverslip. Cortical responses were always recorded through the intact skull. During the experiment isoflurane inhalation anesthesia was applied through a plastic mask and maintained at 0.5-0.6%.

#### Mouse preparation for repeated imaging experiments

Repeated intrinsic imaging in the same mice was performed as previously described (Kaneko et al., 2008; Teichert et al., 2018a). Briefly, after the first imaging (0 days) session the skin was re-sutured and animals were returned to their standard cages. During the subsequent days animals received a daily injection of carprofen (5 mg/kg, s.c.) for pain prevention. Before the next imaging session (after 7 days) the previously closed eye and the skin covering the visual cortex was re-opened and imaging was performed as described above (**Figure 1**).

**Figure 1:**
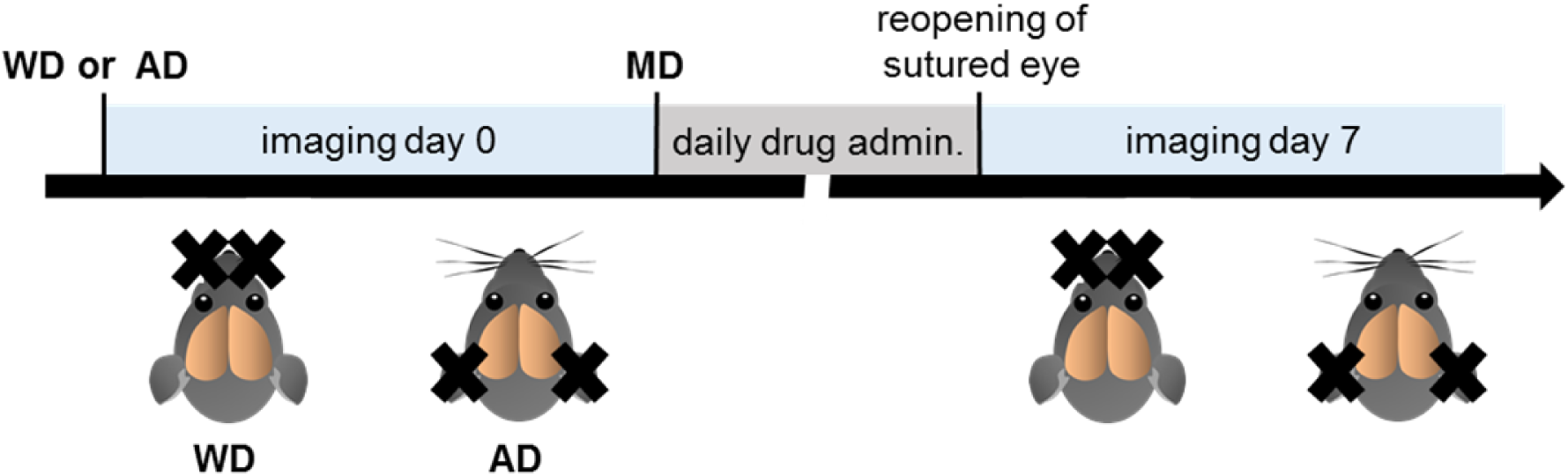
Experimental time course. WD or AD respectively were always performed before the first imaging session (day 0). Subsequently after the initial imaging session animals were monocularly deprived. During the following days the received daily drug infusions (saline, diazepam, CPP, respectively, or a mixture of CPP and diazepam). The final imaging experiment was performed after 7 days.

#### Imaging of visual cortex

Responses of mouse primary visual cortex were recorded as described previously (Teichert et al., 2018a). Briefly, the method uses a periodic stimulus that is presented to the animal for some time and cortical responses are extracted by Fourier analysis. In our case, the visual stimulus was a drifting horizontal light bar of 2° width, 100% contrast and with a temporal frequency of 0.125 Hz. The stimulus was presented on a high refresh rate monitor (Hitachi Accuvue HM 4921-D) placed 25 cm in front of the animal. Visual stimulation was adjusted so that it only appeared in the binocular visual field of the recorded hemisphere (−5° to +15° azimuth, −17° to +60° elevation). The stimulus was presented to the contra or ipsilateral, respectively for 5 min and was repeated for about 35 times during one presentation period.

#### CCD camera recording procedure

Using a Dalsa 1M30 CCD camera (Dalsa, Waterloo, Canada) with a 135×50 mm tandem lens (Nikon, Inc., Melville, NY), we first recorded images of the surface vascular pattern via illumination with green light (550±2 nm) and, after focusing 600 µm below the pial surface, intrinsic signals were obtained via illumination with red light (610±2 nm). Frames were acquired at a rate of 30 Hz and temporally averaged to 7.5 Hz. The 1024×1024 pixel images were spatially averaged to a 512×512 resolution. We always imaged the left hemisphere of the animals.

#### Data analysis

From the recorded frames the signal was extracted by Fourier analysis at the stimulation frequency and converted into amplitude and phase maps using custom software (Kalatsky and Stryker, 2003). In detail, from a pair of the upward and downward maps, a map with absolute retinotopy and an average magnitude map were computed. For data analysis we used the MATLAB standard as described previously (Cang et al., 2005; Lehmann and Lowel, 2008). The magnitude component represents the activation intensity of the visual cortex. Since high levels of neuronal activity decrease oxygen levels supplied by hemoglobin and since deoxyhemoglobin absorbs more red light (610±2 nm), the reflected light intensity decreases in active cortical regions. Because the reflectance changes are very small (less than 0.1%) all amplitudes are multiplied with 10^4^ so that they can be presented as small positive numbers. Thus, the obtained values are dimensionless. Amplitude maps were obtained by averaging the response amplitudes of individual pixels from maps to upward and downward moving bars. The ocular dominance index (ODI) was computed as (C-I)/(C+I) with C and I representing the peak response amplitudes of V1 elicited by contralateral eye and ipsilateral eye stimulation, as described previously (Cang et al., 2005; Kaneko et al., 2008). To each condition we took at least three magnitudes of V1 responsiveness and averaged them for data presentation.

### Whisker deprivation (WD) and auditory deprivation (AD)

WD (n=11) and AD (n=9) for imaging experiments were always performed before the first imaging session (**Figure 1**) (Teichert et al., 2018a). WD was performed as described previously (He et al., 2012; Teichert et al., 2018a). Briefly, animals were deeply anesthetized with 2% isoflurane in a mixture of 1:1 O_2_/N_2_O applied through a plastic mask. The eyes of the animal were protected with silicon oil. Whiskers (macro vibrissae) were plucked bilaterally using fine forceps. Subsequently, mice received an injection of carprofene (4 mg/kg, s.c.) for pain prevention and were either imaged or returned to their standard cages for 7 days. Over the following days whiskers were re-shaved every other day, and animals received a daily administration of carprofene (4 mg/kg, s.c.). Control animals (for HPLC experiments) were sham plucked under the same anesthesia regime by gently pulling on each whisker but leaving them intact as described previously (Teichert et al., 2018b). Control mice also received carprofene injections (4 mg/kg, s.c.). AD was always induced by bilateral malleus removal as described previously (Teichert and Bolz, 2017b, a; Teichert et al., 2017). Briefly, animals were deeply anesthetized with 2% isoflurane in a mixture of 1:1 O_2_/N_2_O applied through a plastic mask. Additionally, mice received a subcutaneous injection of carprofene (4 mg/kg, s.c.) for pain prevention. The eyes of the animal were protected with silicon oil. The tympanic membrane was punctured and the malleus was removed under visual control through this opening using fine sterilized forceps. Great care was taken to avoid any destruction of the stapes and the oval window which is visible through the hearing canal (see (Tucci et al., 1999)). Over the following days animals received a daily administration of carprofene (4 mg/kg, s.c.).

### Monocular deprivation (MD)

MD was always performed after the first imaging session in a total of 26 mice (**Figure 1**) (Teichert et al., 2018a). For this, we increased the isoflurane concentration to 2% in a mixture of 1:1 N_2_O and O_2_. Lid margins of the right eye were trimmed and an antibiotic ointment was applied. Subsequently the right eye was sutured. After MD animals received one injection of carprofene (4 mg/kg, s.c.) and were returned to their standard cages. All animals were checked daily to ensure that the sutured eye remained closed during the MD time. Over the following days animals received a daily administration of carprofene (4 mg/kg, s.c.).

### Saline, diazepam and CPP injections

For repeated optical imaging recordings mice of different experimental groups received daily injections of different drugs. The first injection was always performed immediately after the first imaging sessions (**Figure 1**). In monocularly deprived control mice in which we also performed WD or AD, respectively, we daily injected saline (i.p., 0.12 ml, n=9). To raise cortical inhibition in monocularly deprived WD and AD mice, we intraperitoneally injected 0.12 ml diazepam solution (in saline, 1mg/kg, i.p., 0.12 ml, n=11) daily. To investigate the effects the effects of pure diazepam treatment in MD mice, animals received a daily injection of a diazepam solution (in saline, 1mg/kg, i.p., 0.12 ml, n=3). Furthermore, to block the N-methyl-D-aspartate (NMDA)-receptor in monocularly deprived WD mice which also received diazepam, we daily injected a mixture containing diazepam (1mg/kg) and (R,S)-3-(2-carbooxypiperazin-4-yl)propyl-1-phosphonic (CPP, Abcam, 12-15 mg/kg, n=3) (Sato and Stryker, 2008; Teichert et al., 2018a) in a volume of 0.12 ml (in saline, i.p.).

### Statistical analysis

The normal distribution of the values in each group was analyzed and confirmed by the Kolmogorov-Smirnov test. In addition, the F-test confirmed the equal distribution of values between groups and, thus, allowed us to compare before and after optical imaging values using a parametric paired *t-*test. HPLC data were analyzed by unpaired t-tests. In the graphs, the levels of significance were set as: *: p<0.05, **: p<0.01, ***: p<0.001. Data were analyzed using GRAPHPAD PRISM 7.0 and are presented as data points of individual animals together with means and standard error of the mean (s.e.m.).

## Results

### WD cross-modally decreases GABA levels in V1

It has been described that a reduction in GABAergic inhibition and, thus, an alteration of the balance between excitation and inhibition (E/I ratio) in V1, plays a pivotal role for the restoration of cortical plasticity (He et al., 2006; Maya-Vetencourt et al., 2008; Harauzov et al., 2010). Since we could previously demonstrate that bilateral WD reinstalled OD plasticity in fully adult mice (Teichert et al., 2018a), we examined whether this cross-sensory plasticity also leads to changes in the E/I ratio in V1. For this, we quantified glutamate and GABA levels in V1 by post-mortem HPLC analysis. While glutamate levels did not change 7 days after WD (n=5) compared to control mice (n=5, control: 100±2.12%, WD: 96.49±3.53%, t(8)=0.852, p=0.42; unpaired t-test, **Figure 2 A**), there was a significant reduction in V1 GABA concentration by about 14% (n=6, control: 100±1.90%, WD: 86.20±3.43%, t(9)=3.313, p=0.009; unpaired t-test, **Figure 2 B**). Hence, the glutamate/GABA ratio was markedly increased 7 days after WD (control: 100.37±1.56, WD: 114.86±1.61, t(8)=0.168, p=0.003; unpaired t-test, **Figure 2 C**). These data suggest that WD cross-modally decreases GABAergic inhibition in the spared V1 and thereby shifts the E/I balance in favor of excitation.

**Figure 2:**
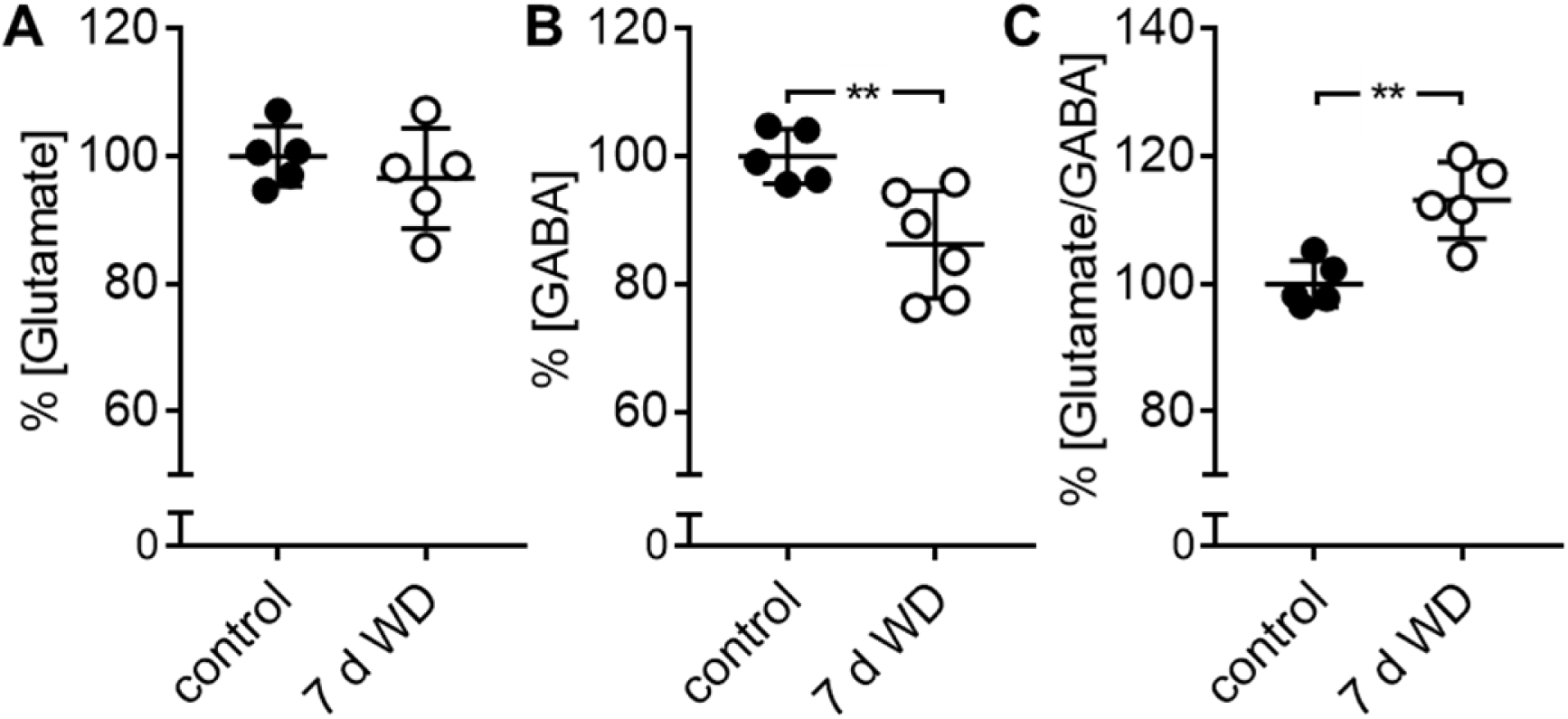
WD cross-modally decreases GABAergic inhibition in V1. **(a)** 7 days of WD did not change V1 glutamate levels (n=5) compared to normal control mice (n=5), as revealed by post-mortem-HPLC analysis of V1 tissue. **(b)** Significant reduction of V1 GABA levels at 7 days after WD (n=6). (c) Hence, there was a significant increase in the glutamate/GABA ratio in WD mice after this time. Open and filled circles represent measurements of individual animals and are presented together with the mean ± s.e.m. **: p<0.01

### WD combined with systemic diazepam administration induces juvenile-like OD plasticity

Many studies could demonstrate that OD plasticity which could be restored by interventions decreasing GABA levels in V1 can be prevented by treatments strengthening GABAergic transmission in the cortex. A common tool used for this purpose is allosteric activation of GABA receptors by diazepam (Sale et al., 2007; Maya-Vetencourt et al., 2008; Spolidoro et al., 2011; Greifzu et al., 2014). If administrated systemically, a dosage of 1 mg/kg has been shown to reliably block OD plasticity (Greifzu et al., 2014; Stodieck et al., 2014). Using repeated intrinsic signal imaging, we measured V1 activity driven by visual stimulation of either the contralateral or ipsilateral eye at 0 and 7 days in whisker and monocularly deprived mice which received daily injections of diazepam (1 mg/kg, i.p., WD+MD+Diaz mice, n=6). In control mice we also combined WD with MD for 7 days but these animals were daily treated with saline (WD+MD+Saline mice, n=5).

**Figure 3 A** (upper part, black frame) depicts representative V1 activity maps elicited by contra or ipsilateral eye stimulation in control mice before (0 days) and after 7 days of WD combined with MD. Generally, darker activity maps indicate higher visually driven V1 responses. While V1 responsiveness to contralateral (closed) eye stimulation did not change during the time tested, there was a marked increase of V1 input strength evoked by visual stimulation of the ipsilateral (open) eye. These data are in correspondence with our recent findings that WD reinstalled “adult-like” OD plasticity in fully adult mice (Teichert et al., 2018a). Surprisingly, enhancing cortical inhibition with diazepam in WD+MD mice did not abolish OD-plasticity as expected, but instead led to a dramatic reduction of V1 responses mediated by the previously closed contralateral eye, whereas ipsilateral eye driven V1 activity remained unchanged after 7 days (**Figure 3 A**, middle part, blue framed). Thus, combining WD and MD together with diazepam administration did also induce OD-plasticity, which was, however, mediated by different V1 input changes than in saline treated WD mice.

**Figure 3:**
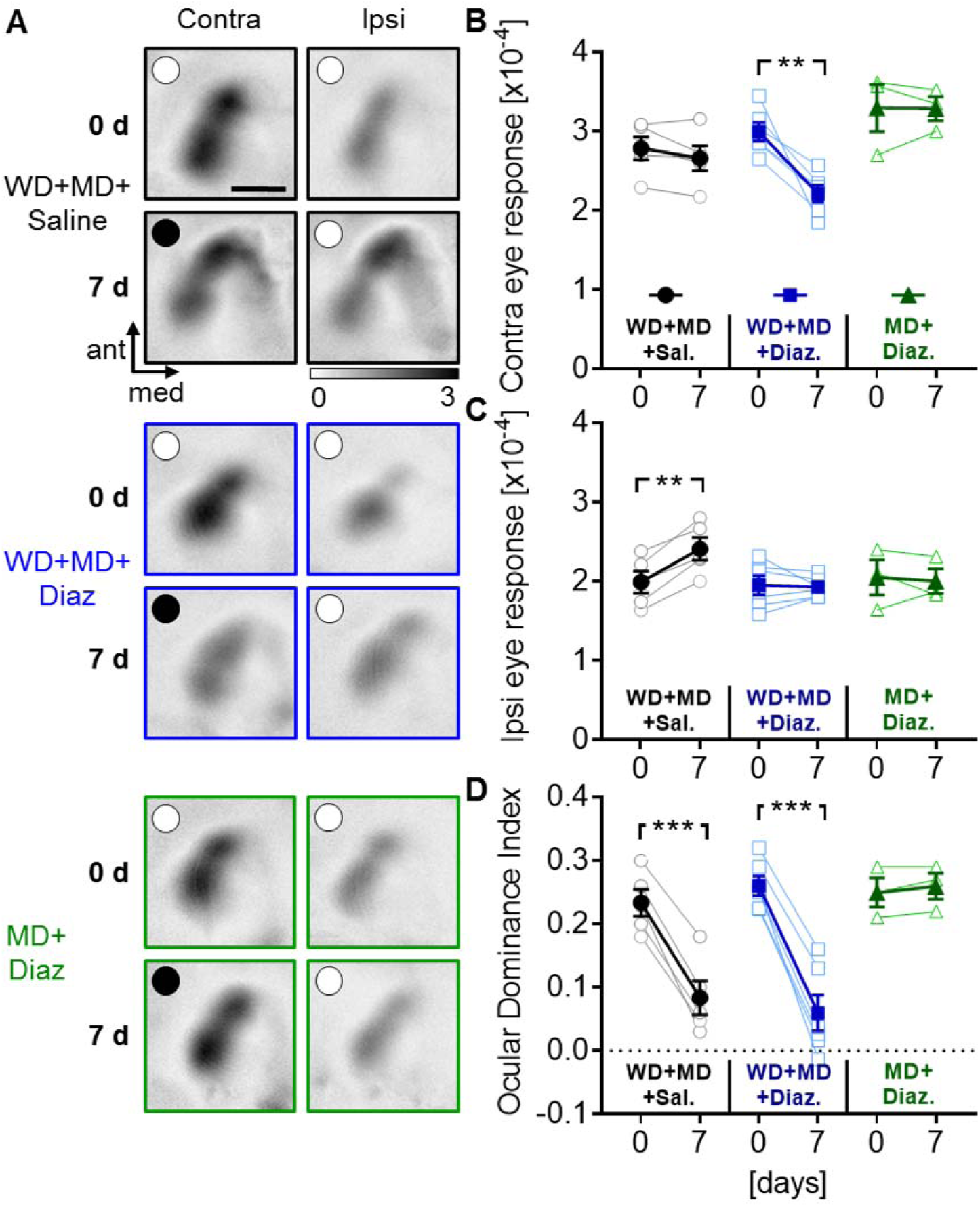
Increasing cortical inhibition by diazepam in WD mice restores “juvenile-like” OD plasticity. **(A)** Representative V1 response maps obtained after visual stimulation of either the contralateral or ipsilateral eye before and after MD. Upper part (black framed): in saline treated control mice (n=5) combined WD and MD for 7 days induced a strengthening of V1 input through the ipsilateral (open) eye, whereas contralateral (closed) eye input to V1 did not change. Middle part (blue framed): daily administrations of diazepam in monocularly deprived WD mice (n=6), however, induced a reduction of V1 responses elicited by contralateral (closed) eye stimulation, whereas ipsilateral (open) eye input did not change. Lower part (green framed): Diazepam treatment of mice which were only monocularly deprived did not affect V1 input strength (n=3). **(B, C)** Quantification of V1 activity changes described above. **(D)** We found significant ODI reductions in control mice treated with saline after WD and MD but also after diazepam treatment. However, in mice which only received a MD, increasing inhibition by diazepam did not affect the ODI. Open circles in A: open eye, closed circles in a: closed eye. In B-D: Open circles, squares and triangles represent measurements of individual animals and filled symbols represent means ± s.e.m. *p<0.05, **p<0.01, ***p<0.001, Scale bar: 1 mm

Quantification of V1 activation showed that contralateral (closed) eye input remained unchanged in saline treated control mice, while ipsilateral (open) eye input significantly increased 7 days after WD and MD (contra: 2.78±0.144 (x10^-4^) vs 2.66±0.16 (x10^-4^), t(4)=1.827 p=0.147; ipsi: 1.99±0.14 (x10^-4^) vs 2.41±0.14 (x10^-4^), t(4)=6.647, p=0.0027, paired t-tests, **Figure 3 B, C**). Thus, the ocular dominance index (ODI) significantly decreased from 0.23±0.02 to 0.08±0.03 (t(4)=10.8, p=0.0004, paired t-test, **Figure 3 D**). However, in diazepam treated monocularly deprived WD mice V1 activity evoked by visual stimulation of the previously closed contralateral eye significantly decreased, whereas ipsilateral (open) eye mediated V1 responses remained unchanged (contra: 3.00±0.11 (x10^-4^) vs 2.22±0.11 (x10^-4^), t(5)=4.603, p=0.0058; ipsi: 1.95±0.12 (x10^-4^) vs 1.93±0.05 (x10^-4^), t(5)=0.257, p=0.808; paired t-tests, **Figure 3 B, C**). Hence, there was a significant reduction of the ODI from 0.26±0.020 to 0.06±0.03 after 7 days (t(5)=12.26, p=6.4×10^-5^, paired t-test, **Figure 3 D**).

The fact that diazepam activates GABA_A_ receptors and reduces the neuronal activity of postsynaptic pyramidal neurons (Salin and Prince, 1996), raises the possibility that diazepam treatment per se induces a general reduction of V1 responsiveness to visual stimuli, which could in turn explain the above mentioned results. To address this issue, we chronically measured V1 activity evoked by contra and ipsilateral eye stimulation in another group of monocularly deprived mice which only received daily diazepam injection. As illustrated in **Figure 3 A** (lower part, green framed), V1 response strength evoked by contra (closed) and ipsilateral (open) eye stimulation remained unchanged after 7 days. Quantification revealed that diazepam treatment alone did not change visually evoked V1 activity elicited by stimulation of the previously closed contralateral and open ipsilateral eye, respectively (contra: 3.30±0.30 (x10^-4^) vs 3.30±0.15 (x10^-4^), t(2)=0.042, p=0.97; ipsi: 2.05±0.22 (x10^-4^) vs 2.00±0.15 (x10^-4^), t(2)=0.296, p=0.97; paired t-tests, **Figure 3 B, C**). Thus, the ODI remained stable in these mice (0.25±0.02 vs 0.26±0.02, t(2)=1.732, p=0.23; paired t-test, **Figure D**). These data indicate that increasing cortical inhibition by diazepam does not decrease visually evoked V1 responses and, does also not reactivate OD plasticity in fully adult mice. Hence, the different V1 input changes in monocularly deprived WD mice and in WD+MD mice which also received diazepam, cannot be explained by the effects of diazepam treatment alone.

OD shifts after MD mediated by a strengthening of V1 input through the ipsilateral (open) eye are characteristically found after MD in young adult mice around postnatal day 60 (Sawtell et al., 2003; Sato and Stryker, 2008; Ranson et al., 2012). Hence, here we confirmed that WD combined with MD (control mice) reinstalls this “adult” form of OD plasticity in fully adult mice, as described previously (Teichert et al., 2018a). However, MD induced OD-shifts which are caused by a reduction of contralateral (closed) eye V1 input are typically present in juvenile mice around postnatal day 28 (Gordon and Stryker, 1996). Thus, our results suggest that the combination of WD, MD and diazepam administration restores “juvenile-like” OD plasticity.

### Increasing inhibition after auditory deprivation also reactivates juvenile plasticity

Recently, we could demonstrate that an auditory deprivation (AD), induced by bilateral malleus removal, cross-modally restores “adult-like” OD plasticity in fully adult mice (Teichert et al., 2018a). Hence, we next investigated whether increasing inhibition by diazepam also changes the signature of OD plasticity form “adult” to “juvenile-like” in AD mice, again using repeated intrinsic signal imaging (n=5, **Figure 4 A**). Another group of mice were monocularly and auditorily deprived and received daily injections of saline (n=4). Quantification of V1 activity revealed that in saline treated mice V1 responses elicited by contralateral eye stimulation remained unchanged whereas there was a strong increase of V1 activity evoked by visual stimulation of the ipsilateral (open) eye (contra: 3.0±0.12 (x10^-4^) vs 2.93±0.08 (x10^-4^), t(3)=0.66, p=0.55; ipsi: 2.05±0.15 (x10^-4^) vs 2.58±0.11 (x10^-4^), t(3)=13.48, p=0.0009; paired t-tests, **Figure 4 B**, **C**). Thus, the average ODI significantly shifted from 0.23±0.01 to 0.09±0.02 (t(3)=12.96, p=0.0009; paired t-test, **Figure 4 D**). These data confirm our previous findings that AD reactivates “adult-like” OD plasticity in fully adult mice. However, increasing inhibition by diazepam in monocularly deprived AD mice induced a loss of V1 input strength through the contralateral (closed) eye, whereas V1 input through the ipsilateral (open) eye did not change (contra: 2.96±0.19 (x10^-4^) vs 2.36±0.19 (x10^-4^), t(4)=12.81, p=0.0002; ipsi: 0.19±0.22 (x10^-4^) vs 2.06±0.20 (x10^-4^), t(4)=1.108, p=0.33; paired t-test, **Figure 4 B, C**). These differential changes of V1 input led to a significant reduction of the ODI in V1 of these mice (0.27±0.03 vs 0.08±0.03, t(4)=10.46, p=0.0005; paired t-test, **Figure 4 D**). These data indicate that increasing cortical inhibition after AD induces “juvenile-like” OD plasticity, like above described for WD mice. Hence, our results suggest that the restoration of “juvenile-like” V1 plasticity after cross-sensory deprivation and increasing inhibition is a general feature of cross-modal plasticity in V1.

**Figure 4:**
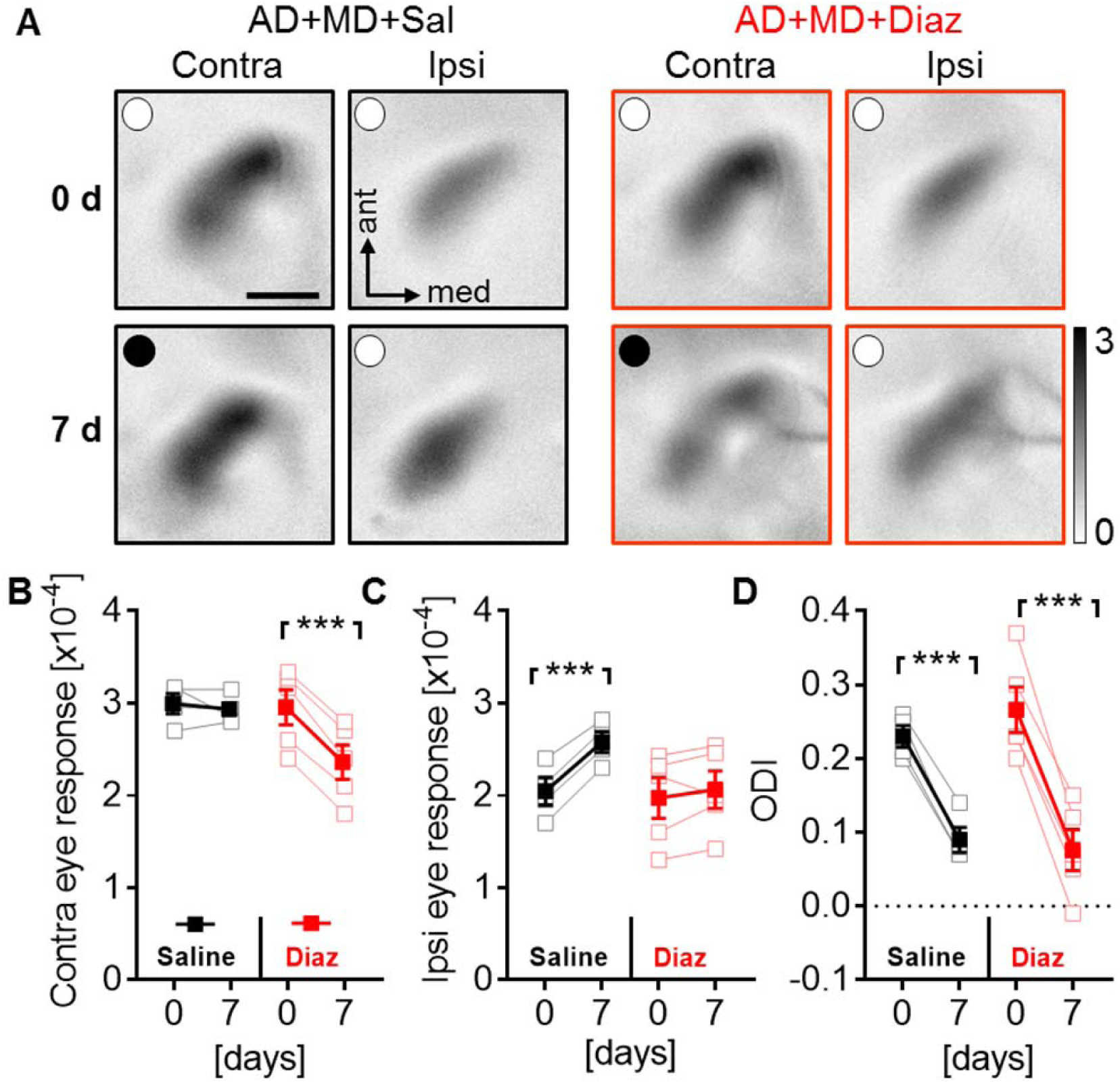
Increasing inhibition by diazepam after AD cross-modally restores “juvenile-like” OD plasticity in fully adult mice. **(A)** Representative amplitude maps obtained after visual stimulation of either the contralateral (previously closed) or ipsilateral (open) eye before and after MD in auditorily deprived mice treated either with saline (n=4) or diazepam (n=5), respectively. (**B**) V1 activity elicited by contralateral eye stimulation remained unchanged in saline but significantly decreased in diazepam treated mice. (**C**) V1 input through the ipsilateral eye significantly increased in mice which received daily saline injections but did not change in mice after diazepam treatment. (**D**) In mice of both groups there was a significant reduction of the ODI. Open circles in A: open eye, closed circles in a: closed eye. In B-D: Open circles, squares and triangles represent measurements of individual animals and filled symbols represent means ± s.e.m. *p<0.05, **p<0.01, ***p<0.001, Scale bar: 1 mm

### OD plasticity after WD and diazepam treatment depends on NMDA receptor activation

We could previously demonstrate that WD induced OD plasticity requires NMDA receptor activation (Teichert et al., 2018a). Hence, we next examined whether this is also the case for OD shifts induced by WD and diazepam treatment. Therefore, we performed repeated optical imaging experiments in monocularly deprived WD mice which daily received a cocktail containing diazepam (1 mg/kg, i.p.) and CPP (15 mg/kg, i.p. WD+MD+Diaz+CPP mice, n=3), a competitive NMDA receptor antagonist (Sato and Stryker, 2008). As shown in **Figure 5 A** this treatment completely prevented V1 activity changes, which occured in monocularly deprived WD mice treated with diazepam alone. Quantification of V1 activation showed that neither contralateral (closed) nor ipsilateral (open) eye input to V1 changed in these mice (contra: 3.06±0.33 (x10^-4^) vs 3.02±0.29 (x10^-4^), t(2)=0.598, p=0.61; ipsi: 2.12±0.16 (x10^-4^) vs 2.12±0.30, t(2)=0.025, p=0.98; paired t-test, **Figure 5 B, C**). As a direct consequence the ODI remained unaltered after 7 days of WD+MD and treatment with diazepam and CPP (0.23±0.03 vs 0.22±0.04, t(2)=0.961, p=0.44, paired t-test, **Figure 5 D**). Taken together, these data suggest that “juvenile-like” OD shifts induced by WD+MD and diazepam treatment are mediated by NMDA receptor activation, too. In general, these results underline the pivotal role of NMDA receptors in cross-modally restored OD plasticity in fully adult mice (Teichert et al., 2018a).

**Figure 5:**
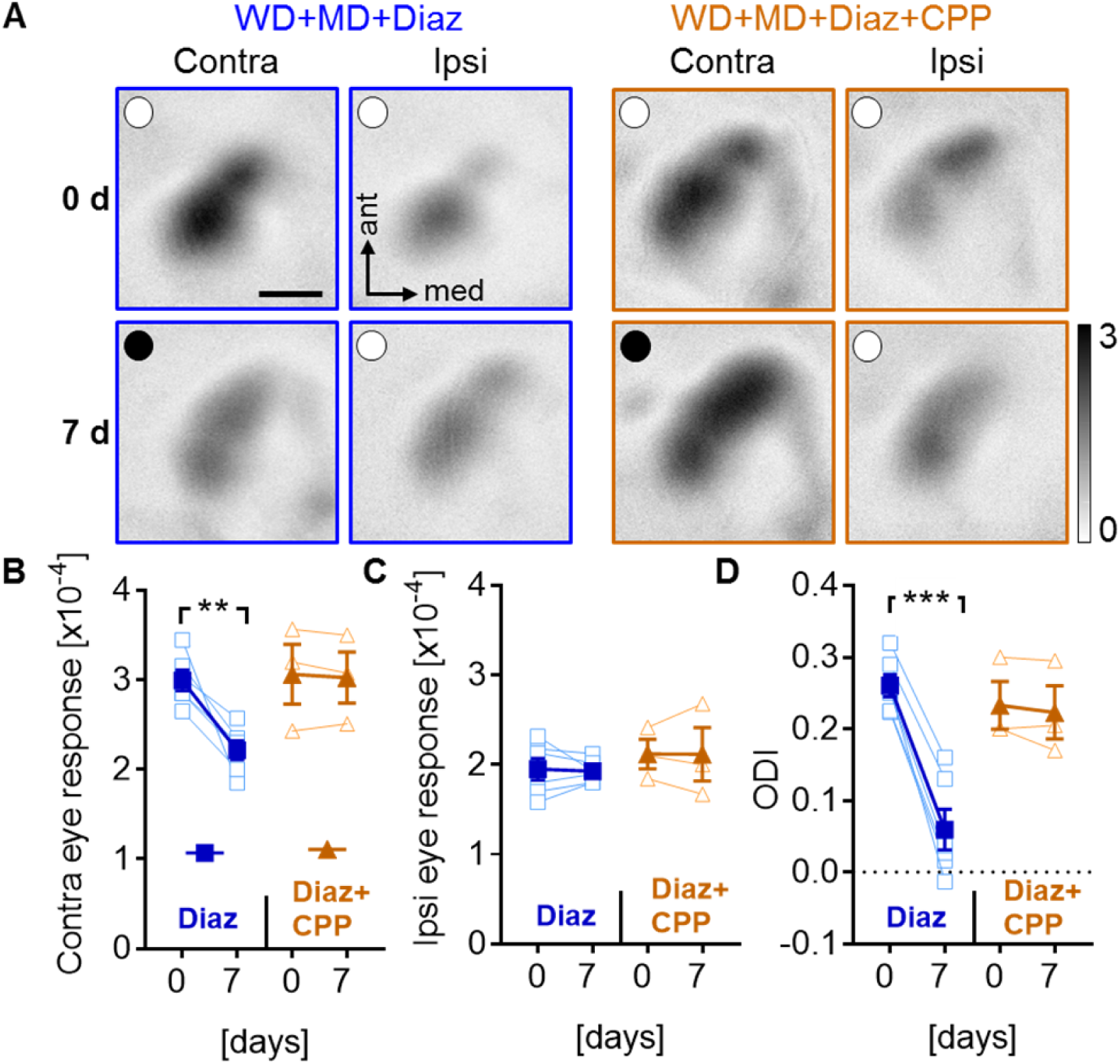
Cross-modally restored “juvenile-like” OD plasticity after diazepam treatment depends on the NMDA receptor. **(A)** Representative V1 response maps evoked by contra (previously closed) or ipsilateral (open) eye stimulation, respectively, before and after MD in WD mice daily treated with diazepam (n=6) or a cocktail containing diazepam and CPP, a competitive NMD receptor blocker (n=3). **(B)** While diazepam treatment in monocularly deprived WD mice decreased V1 responsiveness to contralateral eye stimulation, additional CPP treatment abolished these changes. **(C)** There were no alteration in the V1 input strength through the ipsilateral eye in mice of both groups. **(D)** Additional CPP treatment prevented OD shifts in diazepam treated mice after AD and MD. Open circles in A: open eye, closed circles in a: closed eye. In B-D: Open circles, squares and triangles represent measurements of individual animals and filled symbols represent means ± s.e.m. *p<0.05, **p<0.01, ***p<0.001, Scale bar: 1 mm

## Discussion

In sensory systems the capacity of the brain to undergo experience dependent plastic changes is typically restricted to well-defined time windows soon after birth (Wiesel and Hubel, 1963; Hensch, 2005). However, after the end of these critical periods, brain plasticity dramatically declines. Because of its potential translational implications, restoring cortical plasticity in adults is a topic of great scientific interest.

We could recently demonstrate that the deprivation of different non-visual sensory modalities reinstalled OD plasticity in V1 of fully adult mice (Teichert et al., 2018a). Specifically, both WD and AD, combined with MD for 7 days induced OD shifts which were mediated by an increased V1 responsiveness after ipsilateral (open) eye stimulation, whereas contralateral (closed) eye input to V1 did not change (Teichert et al., 2018a). This is the characteristic signature of “adult-like” plasticity, typically found in young adult mice around PD 60 (**Figure 6**) (Sato and Stryker, 2008; Ranson et al., 2012). However, here we made the surprising finding that enhancing GABAergic inhibition did not prevent these OD shifts, as suggested by many previous studies (**Figure 6**) (Sale et al., 2007; Maya-Vetencourt et al., 2008; Greifzu et al., 2014; Stodieck et al., 2014). Instead, after combining either WD or AD, with diazepam injections, OD shifts were still present but were now mediated by a reduction in contralateral (closed) eye input to V1 but the ipsilateral (open) eye remained unaltered (**Figure 3, 4**), which is the characteristic feature of “juvenile-like” OD plasticity, generally present in very young mice around 30 days of age (**Figure 6**) (Gordon and Stryker, 1996; Ranson et al., 2012). These results indicate that mechanisms, which mediates cross-modally induced OD shifts, change after artificially increasing inhibition in the cortex. Thus, our results suggest that the alteration of GABAergic inhibition in V1 is not the only mechanism involved in mediating cross-modally restored V1 plasticity.

**Figure 6:**
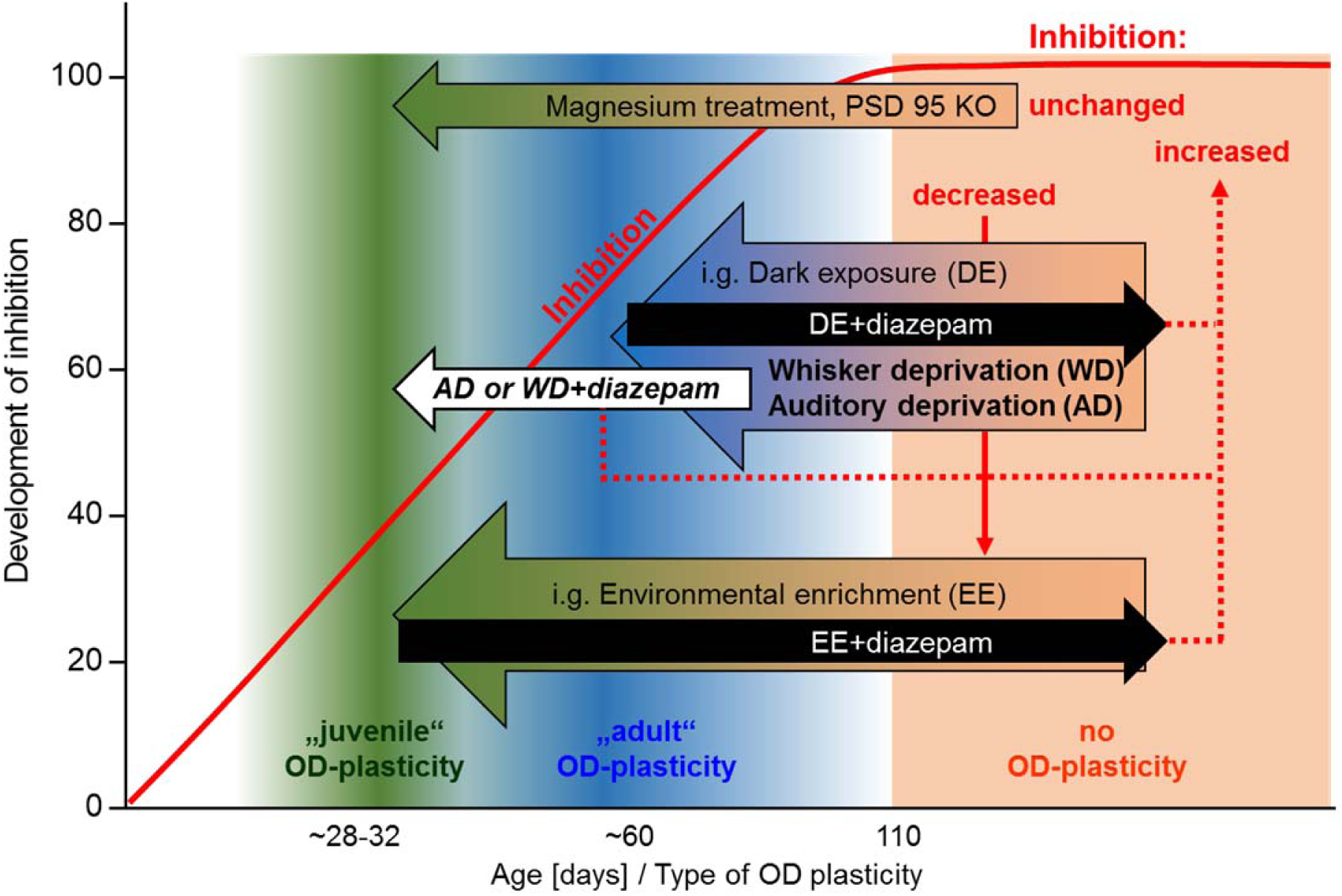
Developmental increase of GABAergic inhibition in V1 is associated by a decline in V1 plasticity. Between 28 and 32 days of age relatively low inhibition levels cause high plasticity levels in mouse V1. MD during this critical period induces a depression of the contralateral (closed) eye input to V1, a signature of “juvenile” OD plasticity (Gordon and Stryker, 1996; Hensch and Fagiolini, 2005; Ranson et al., 2012). In young adult mice, at around 60 days of age, the quality of OD shifts changes, as MD causes potentiation of ipsilateral (open) eye input to V1, the characteristic feature of “adult” OD plasticity (Sato and Stryker, 2008; Ranson et al., 2012). In mice older than 110 days, when cortical inhibition is high, OD plasticity is completely absent (Lehmann and Lowel, 2008). However, interventions that decrease GABAergic inhibition in V1, such as dark exposure (DE) (He et al., 2006; Stodieck et al., 2014) or environmental enrichment (EE) (Sale et al., 2007; Greifzu et al., 2014), can restore “adult-like” and “juvenile-like” OD plasticity. Increasing GABAergic inhibition by diazepam, however, prevents OD plasticity after these interventions (Greifzu et al., 2014; Stodieck et al., 2014). We could previously demonstrate that whisker deprivation (WD) reinstalled the “adult-like” form of OD plasticity in fully adult mice older than 110 days (Teichert et al., 2018a). Here, we show that WD decreases GABA levels in V1, suggesting that this alteration is causal for the restoration of “adult-like” plasticity”. However, as shown here, increasing GABAergic inhibition by diazepam, which should shift the impact of cortical GABA to the “no-OD-plasticity-stage” did not prevent OD shifts. Strikingly, the signature of OD plasticity changed from “adult” to “juvenile”. Interestingly, this was also the case after diazepam treatment of AD mice. Thus, the cross-modal restoration of “juvenile-like” plasticity in V1 does not solely depend on alterations of the inhibitory tone, but also requires alternative, so far unknown cellular and molecular mechanisms.

To the best of our knowledge, this finding is unique. Previous interventions such as dark exposure (He et al., 2006; Stodieck et al., 2014), environmental enrichment (Sale et al., 2007; Baroncelli et al., 2010; Greifzu et al., 2014), food restriction (Spolidoro et al., 2011) and fluoxetine administrations (Maya-Vetencourt et al., 2008), all restored OD plasticity in V1 of fully adult mice. A common thread of these treatments is that they all cause lower levels of GABAergic inhibition in V1. However, while dark exposure restored the “adult” form of OD plasticity (Stodieck et al., 2014), environmental enrichment and fluoxetine treatment for example reinstalled the “juvenile” form (**Figure 6**) (Maya-Vetencourt et al., 2008; Greifzu et al., 2014). Despite these differences, mechanistically, the decrease in inhibition appears to be the central hub, because increasing inhibition by diazepam prevents OD shifts after these interventions (**Figure 6**) (Sale et al., 2007; Maya-Vetencourt et al., 2008; Spolidoro et al., 2011; Greifzu et al., 2014; Stodieck et al., 2014).

This idea is further supported by the finding that OD plasticity in fully adult rodents can also be restored by reducing GABA synthesis or antagonizing GABA receptors in the adult V1 (Harauzov et al., 2010). Indeed, in accordance with these findings, we show here that cross-modally induced OD plasticity is also associated with a decrease of V1 GABA levels (**Figure 2**). Notably, Harauzov and colleagues (2010) could also demonstrate that reducing the inhibitory tone facilitated white matter stimulation induced long-term potentiation (LTP) in V1 slices, whereas the thresholds for LTD remained unchanged. They therefore proposed that this treatment would most likely induce “adult-like” OD plasticity after MD (Harauzov et al., 2010), since this type of V1 plasticity is indeed mediated by LTP-like mechanisms (Sawtell et al., 2003; Cooke and Bear, 2014). This idea is also in line with the present finding that both WD and AD, restored the “adult” form of OD plasticity (**Figure 3, 4**). At first glance, these results seem to indicate that the reduction of GABA levels in V1 is causal for the cross-modal restoration of OD plasticity.

Most of the previous studies, which demonstrated that a reduction of cortical GABA levels restores OD plasticity, used *in vivo* brain microdialysis to obtain cortical probes (Sale et al., 2007; Maya-Vetencourt et al., 2008; Baroncelli et al., 2010). The GABA content of these probes was subsequently determined by HPLC analysis. This experimental design enables measuring extracellular GABA levels (Sale et al., 2007; Maya-Vetencourt et al., 2008; Baroncelli et al., 2010). Hence, the GABA levels measured this most likely represent the biologically active GABA in the synaptic clefts. In the present study, we determined GABA levels in micro punches of V1 tissue, which contain both extracellular and intracellular GABA. Thus, we cannot make definite statements about to which amount the extracellular GABA concentration decreased after WD. However, as we found a general cross-modal reduction in V1 GABA levels (**Figure 2**), we believe that this is accompanied with a general reduction of inhibition, as suggested previously (Harauzov et al., 2010; Spolidoro et al., 2011)

Enhancing GABAergic transmission with diazepam did not abolish cross-modally induced OD shifts but instead changed the signature of OD plasticity (**Figure 3, 4**). Hence, our data suggest that the reduction of GABA levels in V1 after WD cannot be the only reason for restoring OD plasticity. Instead, there must be additional mechanisms which mediate cross-modal plasticity. In other words, the WD induced decrease of GABA levels sets V1 of fully adult mice back into a plastic stage facilitating the induction of “adult-like” OD plasticity (**Figure 6**). However, increasing GABAergic inhibition in V1 by diazepam activates other, so far unknown, mechanisms, which do not only require reduced GABA levels in V1, to mediate cross-modally induced “juvenile-like” plasticity. Such alternative mechanisms have been already described. For instance, magnesium treatment, which increased the expression of specific subunits of the NMDA receptor, has been shown to restore “juvenile-like” OD plasticity in fully adult mice without changing the inhibitory tone in V1 (Liu et al., 2015). Likewise, PSD-95 KO mice display a lifelong preservation of “juvenile-like” OD plasticity, which is not accompanied by a reduction of inhibition in V1, and, hence, cannot be prevented by diazepam administration (Huang et al., 2015). Again, these findings demonstrate that there are, indeed, mechanisms which can restore “juvenile-like” plasticity without reducing cortical inhibition in V1 (**Figure 6**).

Generally, in accordance with our previous findings that both WD and AD reactivate “adult-like” OD plasticity in V1 of fully adult mice, we could here demonstrate that the same sensory deprivations (WD and AD) can also reactivate “juvenile-like” plasticity in V1 after increasing inhibition. Collectively, these results suggest a high similarity of mechanisms taking place in V1 after depriving non-visual sensory modalities. However, the cross-modal interplay of primary sensory regions in normal mice appears to be asymmetric (Iurilli et al., 2012; Teichert and Bolz, 2017b). For instance, while sensory evoked activity in the primary somatosensory (S1) and primary auditory cortex (A1) suppresses V1 responses (Iurilli et al., 2012; Teichert and Bolz, 2017b, a), V1 activity evokes depolarizations in S1 and has only little impact on A1 responses (Iurilli et al., 2012). Thus, it might be that deprivation of vision provokes different effects on the remaining sensory cortices such as S1 and A1.

There is increasing evidence that the “juvenile” form of OD plasticity is mediated by LTD-like mechanisms that decrease contralateral (closed) eye input (Kirkwood and Bear, 1994; Heynen et al., 2003; Espinosa and Stryker, 2012; Cooke and Bear, 2014), whereas the “adult” form occurs via LTP-like changes that lead to an increase of open eye input to V1 (Sawtell et al., 2003; Sato and Stryker, 2008; Ranson et al., 2012; Cooke and Bear, 2014). This idea is supported by the finding that both “juvenile” and “adult” plasticity require NMDA receptor activation (Sato and Stryker, 2008). In accordance with these findings we could recently show that cross-modally restored “adult” OD plasticity depends on NMDA receptor activation (Teichert et al., 2018a). As demonstrated in the present study, this is also the case when “juvenile” OD plasticity is induced by diazepam administration in WD mice (**Figure 5**). Thus, these results suggest that diazepam administration in monocularly and whisker deprived mice, facilitates LTD-like mechanisms, whereas MD combined with WD alone facilitates LTP-like changes in V1. Generally, these results also demonstrate the central role of the NMDA receptor in cross-modal plasticity (Teichert et al., 2018a).

### Conclusion

Here we provide evidence that cross-modally induced OD plasticity in fully adult mice older than 110 days (Teichert et al., 2018a) is accompanied by a reduction of GABA levels in the spared V1. This finding is in line with the current view that a decrease in the inhibitory tone is the central hub which mediates restoration of cortical plasticity. In contrast and surprisingly, our results also suggest that mechanisms other than reduced GABA levels mediate the cross-modal restoration of OD plasticity, as increasing the inhibitory tone did not abolish OD plasticity. However, the signature of OD plasticity changed from “adult-like” to “juvenile-like”. While further research is needed to unravel the precise underlying molecular mechanisms, the present results emphasize the power of cross-modally induced plasticity to re-open a window of high plasticity in the fully adult cortex far beyond any sensory critical period. More general, while therapeutic interventions after sensory damage concentrate on the affected sensory modality, there might be a unique opportunity to sharpen and refine the spared senses and thereby partially compensate sensory deficits.

## Conflict of Interest Statement

The authors declare that the research was conducted in the absence of any commercial or financial relationships that could be construed as a potential conflict of interest.

## Funding

This research did not receive any specific grant from funding agencies in the public, commercial, or not-for-profit sectors.

## Acknowledgments

Thanks are due to Elisabeth Meier for excellent technical assistance and Sandra Eisenberg for animal care.

## Author contributions

MT, MI and JB designed the study

MT, MI, FW, CW performed the experiments

MT, MI, and JB analyzed data

MT and JB wrote the paper

## Abbreviations

A1: Primary auditory cortex
AD: Auditory deprivation
CPP: (R,S)-3-(2-carbooxypiperazin-4-yl)propyl-1-phosphonic
E/I: Excitation/Inhibition
GABA: gamma-Aminobutyric acid
HPLC: high performance liquid chromatography
MD: Monocular deprivation
NMDAR: N-methyl-D-aspartate receptor
OD: Ocular dominance
ODI: Ocular dominance index
PD: Postnatal day
PSD-95: postsynaptic density protein 95
S1: Primary somatosensory cortex
V1: Primary visual cortex
WD: Whisker deprivation

## References

Baroncelli L, Sale A, Viegi A, Maya Vetencourt JF, De Pasquale R, Baldini S, Maffei L (2010) Experience-dependent reactivation of ocular dominance plasticity in the adult visual cortex. Exp Neurol 226:100–109.

Bavelier D, Neville HJ (2002) Cross-modal plasticity: where and how? Nature reviews Neuroscience 3:443–452.

Cang J, Kalatsky VA, Lowel S, Stryker MP (2005) Optical imaging of the intrinsic signal as a measure of cortical plasticity in the mouse. Visual neuroscience 22:685–691.

Cooke SF, Bear MF (2014) How the mechanisms of long-term synaptic potentiation and depression serve experience-dependent plasticity in primary visual cortex (vol 369, 20130284, 2013). Philos T R Soc B 369.

Drager UC (1975) Receptive fields of single cells and topography in mouse visual cortex. The Journal of comparative neurology 160:269–290.

Drager UC (1978) Observations on monocular deprivation in mice. Journal of neurophysiology 41:28–42.

Espinosa JS, Stryker MP (2012) Development and plasticity of the primary visual cortex. Neuron 75:230–249.

Gordon JA, Stryker MP (1996) Experience-dependent plasticity of binocular responses in the primary visual cortex of the mouse. The Journal of neuroscience 16:3274–3286.

Greifzu F, Pielecka-Fortuna J, Kalogeraki E, Krempler K, Favaro PD, Schluter OM, Lowel S (2014) Environmental enrichment extends ocular dominance plasticity into adulthood and protects from stroke-induced impairments of plasticity. Proceedings of the National Academy of Sciences of the United States of America 111:1150–1155.

Harauzov A, Spolidoro M, DiCristo G, De Pasquale R, Cancedda L, Pizzorusso T, Viegi A, Berardi N, Maffei L (2010) Reducing Intracortical Inhibition in the Adult Visual Cortex Promotes Ocular Dominance Plasticity. Journal of Neuroscience 30:361–371.

He HY, Hodos W, Quinlan EM (2006) Visual deprivation reactivates rapid ocular dominance plasticity in adult visual cortex. The Journal of neuroscience 26:2951–2955.

He K, Petrus E, Gammon N, Lee HK (2012) Distinct sensory requirements for unimodal and cross-modal homeostatic synaptic plasticity. The Journal of neuroscience 32:8469–8474.

Hensch TK (2005) Critical period plasticity in local cortical circuits. Nature reviews Neuroscience 6:877–888.

Hensch TK, Fagiolini M (2005) Excitatory-inhibitory balance and critical period plasticity in developing visual cortex. Progress in brain research 147:115–124.

Heynen AJ, Yoon BJ, Liu CH, Chung HJ, Huganir RL, Bear MF (2003) Molecular mechanism for loss of visual cortical responsiveness following brief monocular deprivation. Nature neuroscience 6:854–862.

Hofer SB, Mrsic-Flogel TD, Bonhoeffer T, Hubener M (2006) Prior experience enhances plasticity in adult visual cortex. Nature neuroscience 9:127–132.

Huang XJ, Stodieck SK, Goetze B, Cui L, Wong MH, Wenzel C, Hosang L, Dong Y, Lowel S, Schluter OM (2015) Progressive maturation of silent synapses governs the duration of a critical period. Proceedings of the National Academy of Sciences of the United States of America 112:E3131–E3140.

Hubener M, Bonhoeffer T (2014) Neuronal plasticity: beyond the critical period. Cell 159:727–737.

Isstas M, Teichert M, Bolz J, Lehmann K (2017) Embryonic interneurons from the medial, but not the caudal ganglionic eminence trigger ocular dominance plasticity in adult mice. Brain structure & function 222:539–547.

Iurilli G, Ghezzi D, Olcese U, Lassi G, Nazzaro C, Tonini R, Tucci V, Benfenati F, Medini P (2012) Sound-driven synaptic inhibition in primary visual cortex. Neuron 73:814–828.

Kalatsky VA, Stryker MP (2003) New paradigm for optical imaging: temporally encoded maps of intrinsic signal. Neuron 38:529–545.

Kaneko M, Stellwagen D, Malenka RC, Stryker MP (2008) Tumor necrosis factor-alpha mediates one component of competitive, experience-dependent plasticity in developing visual cortex. Neuron 58:673–680.

Kirkwood A, Bear MF (1994) Homosynaptic long-term depression in the visual cortex. The Journal of neuroscience: the official journal of the Society for Neuroscience 14:3404–3412.

Lee HK, Whitt JL (2015) Cross-modal synaptic plasticity in adult primary sensory cortices. Current opinion in neurobiology 35:119–126.

Lehmann K, Lowel S (2008) Age-dependent ocular dominance plasticity in adult mice. Plos One 3:e3120.

Levelt CN, Hubener M (2012) Critical-period plasticity in the visual cortex. Annual review of neuroscience 35:309–330.

Liu HX, Li Y, Wang Y, Wang XX, An X, Wang SY, Chen L, Liu GS, Yang YP (2015) The distinct role of NR2B subunit in the enhancement of visual plasticity in adulthood. Mol Brain 8.

Maya-Vetencourt JF, Sale A, Viegi A, Baroncelli L, De Pasquale R, O’Leary OF, Castren E, Maffei L (2008) The antidepressant fluoxetine restores plasticity in the adult visual cortex. Science 320:385–388.

Ranson A, Cheetham CE, Fox K, Sengpiel F (2012) Homeostatic plasticity mechanisms are required for juvenile, but not adult, ocular dominance plasticity. Proceedings of the National Academy of Sciences of the United States of America 109:1311–1316.

Sale A, Maya Vetencourt JF, Medini P, Cenni MC, Baroncelli L, De Pasquale R, Maffei L (2007) Environmental enrichment in adulthood promotes amblyopia recovery through a reduction of intracortical inhibition. Nature neuroscience 10:679–681.

Salin PA, Prince DA (1996) Spontaneous GABAA receptor-mediated inhibitory currents in adult rat somatosensory cortex. Journal of neurophysiology 75:1573–1588.

Sato M, Stryker MP (2008) Distinctive features of adult ocular dominance plasticity. The Journal of neuroscience: the official journal of the Society for Neuroscience 28:10278–10286.

Sawtell NB, Frenkel MY, Philpot BD, Nakazawa K, Tonegawa S, Bear MF (2003) NMDA receptor-dependent ocular dominance plasticity in adult visual cortex. Neuron 38:977–985.

Spolidoro M, Baroncelli L, Putignano E, Maya-Vetencourt JF, Viegi A, Maffei L (2011) Food restriction enhances visual cortex plasticity in adulthood. Nature communications 2:320.

Stodieck SK, Greifzu F, Goetze B, Schmidt KF, Lowel S (2014) Brief dark exposure restored ocular dominance plasticity in aging mice and after a cortical stroke. Experimental gerontology 60:1–11.

Teichert M, Bolz J (2017a) Data on the effect of conductive hearing loss on auditory and visual cortex activity revealed by intrinsic signal imaging. Data in Brief 14: 659–664.

Teichert M, Bolz J (2017b) Simultaneous intrinsic signal imaging of auditory and visual cortex reveals profound effects of acute hearing loss on visual processing. NeuroImage 159:459–472.

Teichert M, Liebmann L, Hubner CA, Bolz J (2017) Homeostatic plasticity and synaptic scaling in the adult mouse auditory cortex. Scientific reports 7:17423.

Teichert M, Isstas M, Zhang Y, Bolz J (2018a) Cross-modal restoration of ocular dominance plasticity in adult mice. The European journal of neuroscience 47:1375–1384.

Teichert M, Isstas M, Wenig S, Setz C, Lehmann K, Bolz J (2018b) Cross-modal refinement of visual performance after brief somatosensory deprivation in adult mice. The European journal of neuroscience 47:184–191.

Tucci DL, Cant NB, Durham D (1999) Conductive hearing loss results in a decrease in central auditory system activity in the young gerbil. Laryngoscope 109:1359–1371.

Wiesel TN, Hubel DH (1963) Single-Cell Responses in Striate Cortex of Kittens Deprived of Vision in 1 Eye. Journal of neurophysiology 26:1003-&.

Winter C, Djodari-Irani A, Sohr R, Morgenstern R, Feldon J, Juckel G, Meyer U (2009) Prenatal immune activation leads to multiple changes in basal neurotransmitter levels in the adult brain: implications for brain disorders of neurodevelopmental origin such as schizophrenia. The international journal of neuropsychopharmacology / official scientific journal of the Collegium Internationale Neuropsychopharmacologicum 12:513–524.

